# MatClassRSA v2 release: A MATLAB toolbox for M/EEG classification, proximity matrix construction, and visualization

**DOI:** 10.1101/2025.11.19.689115

**Authors:** Bernard C. Wang, Raymond Gifford, Nathan C. L. Kong, Feng Ruan, Anthony M. Norcia, Blair Kaneshiro

## Abstract

MatClassRSA is a MATLAB toolbox that performs magnetoencephalography and electroencephalography (M/EEG) classification and other analyses related to Representational Similarity Analysis (RSA). The toolbox is designed for cognitive neuroscience researchers who wish to perform classification-based decoding analyses of their data—often repeated trials of evoked responses—or derive Representational Dissimilarity Matrices (RDMs) as input to RSA. Organized as a collection of user-called functions, MatClassRSA v2 comprises five main modules: Preprocessing, Reliability, Classification, RDM Computation, and Visualization. The Classification module supports multiple classifiers (LDA, RF, SVM) and classification schemes (e.g., multiclass, pairwise, cross-validated, train-test, hyperparameter optimization) and offers basic statistical analyses via permutation testing. Functions in other modules include, for example, trial-averaging and noise normalization; reliability estimation; non-classification RDM construction; and hierarchical and non-hierarchical clustering visualizations. The toolbox is freely available on GitHub under an MIT license and includes the main codebase, a User Manual, example function calls, and illustrative analyses. This preprint provides a general background of M/EEG classification for RSA as well as a narrative overview of the v2 MatClassRSA release, its updated functionalities, and illustrative analyses performed using the toolbox.

## Introduction

Classification involves the construction of a statistical model from categorically labeled data observations, which is then used to predict labels of new observations [1]. In the case of magnetoencephalography (MEG) or electroencephalography (EEG) data, observations can be single or group-averaged trials, and categorical labels can be, for example, descriptors of stimuli (e.g., ‘face’, ‘cat’, expected, unexpected), tasks (e.g., difficulty, attentional target), or participants (e.g., expert versus novice). M/EEG classification is considered a *decoding* approach, in that ‘stimulus features’ (broadly defined, and encompassing the above labeling examples) are predicted from the brain data [2]. Classification offers several advantages for analyzing multivariate neuroimaging data, including data-driven selection and weighting of features, predictive capabilities, and the ability to provide interpretable results across large stimulus sets.

Both MEG and EEG are considered suitable data modalities for multivariate analyses, including classification [3]. EEG classification has a long history in brain-computer interfaces (BCI), where accurate decoding—often from single data trials—is crucial for system performance [4]. In recent decades, however, M/EEG classification has also gained traction in cognitive neuroscience research and has been used to study, for example, behavioral performance [5] as well as representation of object categories [6–11], language categories [12], and stimulus modality [13]. In these contexts, research goals may shift away from optimizing decoding accuracy, and classification may rather be used to identify salient data features or as a means of obtaining neural measures of distance or similarity across a stimulus set through classifier accuracies or confusions, respectively.

This latter treatment of classifier accuracies as neural proximities supports the use of M/EEG data in Representational Similarity Analysis (RSA). As explained in seminal RSA papers by Kriegeskorte and colleagues [14, 15], the core tenet of this approach is the abstraction of multiple models, response data, or other representations of a stimulus set away from their native formats (e.g., M/EEG trials, image statistics, perceptual ratings) to a Representational Dissimilarity Matrix (RDM) for each, which summarizes all pairwise distances across the set. There exist various methods for computing RDMs, including correlations, difference vectors, or—as noted above—classifier outputs, and pairs of RDMs can be rank-correlated to assess their representational similarity. In all, RDM representations enable data that may have been difficult to compare directly in disparate original modalities—for example, a set of speech waveforms with a set of EEG responses—to be quantitatively compared in a common similarity space.

RSA has been applied in numerous M/EEG studies. Classifier accuracies and confusions have been compared to perceptual, acoustical, descriptor-based, and computational representations of stimuli [10, 12, 16, 17]. Classification-based RDMs can be visualized not only as matrices but also via hierarchical or non-hierarchical clustering methods to convey the neural proximity structure of the stimuli [7, 9, 18]. Moreover, classifications can be performed over temporal and/or spatial subsets of multichannel M/EEG data [6, 7, 9, 19, 20] to highlight spatiotemporal features relevant to discriminating between classes. The high temporal resolution of M/EEG data also makes their RDMs a useful complement to fMRI in multimodal RSA analyses, for example to relate temporal dynamics (from M/EEG RDMs) to precise localization (from fMRI RDMs) by identifying similar RDMs across response modalities [8, 21].

In this preprint, we introduce the v2 release of MatClassRSA, a MATLAB toolbox centered around M/EEG classification for RSA analyses. As summarized in Fig. 1, the toolbox encompasses Preprocessing, Reliability, Classification, RDM Computation, and Visualization functionalities in a modular structure. MatClassRSA is geared toward cognitive neuroscience researchers working with M/EEG data and MATLAB—particularly those working in ERP paradigms, which are a common data type for classification. As MatClassRSA is a function-based toolbox (i.e., there is no graphical user interface (GUI) at this time), users should be comfortable writing code to load input data, call functions, and save output data and figures. In terms of scope, MatClassRSA is designed to work with data that have already undergone standard cleaning steps, and ends at the construction and visualization of RDMs. Users are encouraged to make use of complementary toolboxes such as Delorme and Makeig‘s EEGLAB [22] for data cleaning and Nili et al.‘s RSA Toolbox [23] for downstream RSA analyses.

**Fig. 1.**
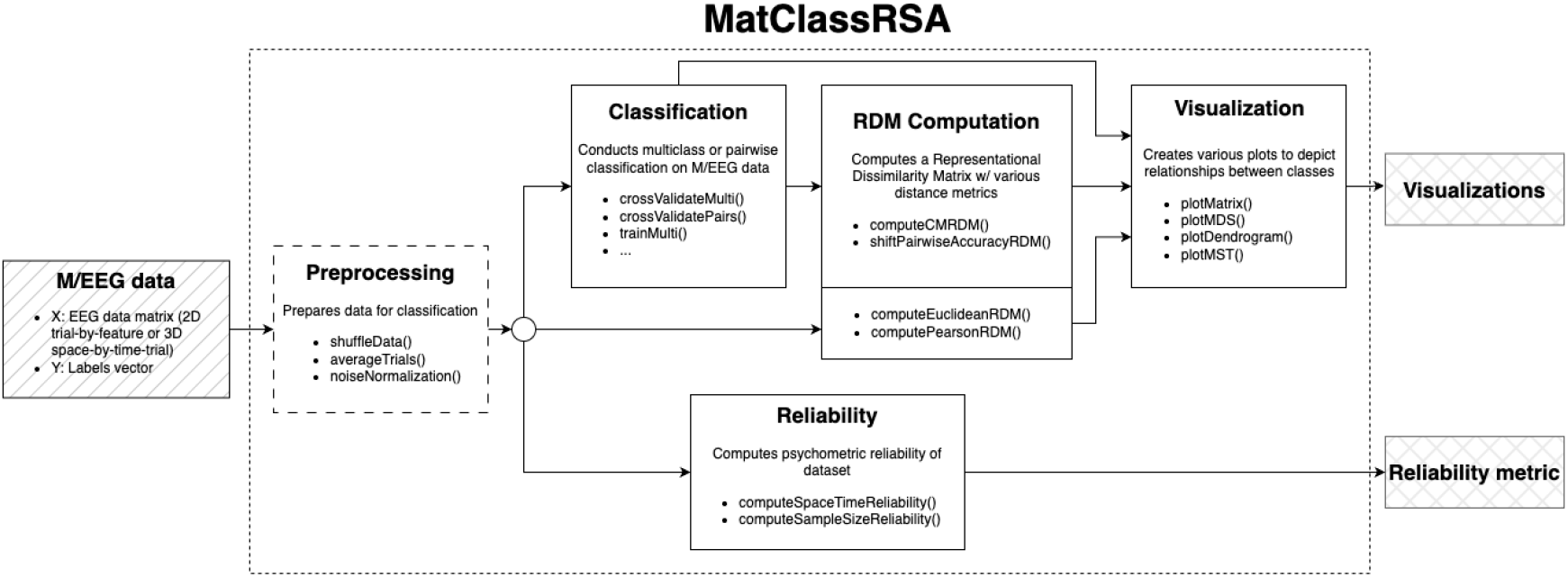
Summary of MatClassRSA functionalities. The toolbox is organized into modules that each focus on different types of analyses. As shown in the flowchart, modules can be arranged according to the user’s analysis goals. MatClassRSA operates on cleaned, ready-to-analyze data matrices and stimulus label vectors.

The MatClassRSA v2 release is a major update and expansion of the original 2017 release [24]. It includes additional functionalities, a restructuring of the codebase, expanded documentation, example function calls, and new illustrative analyses. The v2 release includes three main deliverables:

- The publicly available GitHub repository,^1^ which includes the codebase, User Manual, example function calls, and illustrative analyses. The current release is also archived on Zenodo with a citeable DOI [25].^2^
- This preprint, which provides a narrative overview of the toolbox.
- A standalone dataset containing example data for the illustrative analyses [26].^3^ The GitHub repository includes code to download these data files into the user’s local copy of the toolbox.

The remainder of this preprint is structured as follows. In the next section, we describe the codebase and its contents. After that, we cover the functionalities of the toolbox, including the main user-called functions. Then, as a demonstration of how MatClassRSA functions can be combined for larger-scale analyses, we present a narrative overview of selected illustrative analyses. We close with a discussion of the toolbox’s affordances and limitations.

### Codebase

#### Code access and license

MatClassRSA is freely available on GitHub (see footnote 1). This preprint corresponds to the v2.0 GitHub release. The GitHub release is also archived as a DOI-linked record on Zenodo (see footnote 2). MatClassRSA v2 is released under the MIT license.^4^ If using any part of the codebase, we ask that you cite this preprint as well as the Zenodo entry [25], and to cite the related dataset if using any of those contents for outside projects [26].

### Dependencies

Usage of MatClassRSA requires a MATLAB^5^ license as well as MATLAB’s Parallel Computing Toolbox^6^ and Statistics and Machine Learning Toolbox.^7^ A number of MatClassRSA illustrative analyses require the MATLAB Image Processing Toolbox.^8^ The Support Vector Machine (SVM) classifications of the toolbox use the LIBSVM library [27],^9^ which is included in the MatClassRSA repository; installation instructions are provided in the User Manual. Otherwise, the toolbox is self-contained, meaning that all helper functions are included in the MatClassRSA GitHub repository, and no other external codebases or toolboxes need to be added to the user’s path.

The current release was developed and comprehensively tested on recent MacOS operating systems (12.* and higher) and is expected to work on Linux and Windows platforms as well. The code was developed with the aim of supporting older versions of MATLAB (2016b and higher, e.g., with older versions of input-parser functionality) and has been validated on MATLAB versions 2021a and higher.

#### Folders and files in the repository

The top level of the repository contains the following files and folders:

##### ExampleData

This folder stores example data required to run the illustrative analyses. Due to file size, the example data files are not directly included in the repository; rather, the illustrative analyses include a script that will download the necessary files into this folder.

##### ExampleFunctionCalls

This folder contains one script per user-called MatClassRSA function. Each script contains one or more runnable examples of the corresponding function.

##### IllustrativeAnalyses

This folder contains the script that downloads the example data into the ***ExampleData*** folder (required in order to run the illustrative analyses) as well as one script per illustrative analysis.

##### src

This folder is the main folder for the MatClassRSA codebase. It contains MATLAB package folders (‘+’ folders) for each of the five MatClassRSA modules as well as for the `+Utils` folder containing the helper functions. The `libsvm-3.24` folder is also a subfolder of ***src***.

##### .gitignore

This file lists any files or folders in the repository that should remain untracked by Git. Of special relevance to MatClassRSA users is that all contents of the ***ExampleData*** folder will be ignored.

##### LICENSE.md

License for the repository (MIT license).

##### MatClassRSA v2 User Manual.pdf

This file is the User Manual of the v2 release. It contains detailed information about getting up and running with the toolbox; comprehensive documentation of the user-called functions; supplemental overview and design rationales for the Classification module; details of the illustrative analyses; and brief documentation of helper functions.

##### README.md

README for the repository.

#### Code/data origins and past usage

Design factors of many MatClassRSA functions were drawn from the analyses reported by Kaneshiro et al. [9]. Certain Preprocessing and RDM Computation functions were adapted from or directly use code from Guggenmos et al. [28];^10^ their specific attribution is provided here below, in the User Manual, and in MatClassRSA function docstrings. The MatClassRSA example data [26] include data files from other published datasets [29, 30], as attributed in the dataset documentation. Previous versions of MatClassRSA functions were used in published works by Kong et al. [10] and Losorelli et al. [18].

### Functionalities of the toolbox

The functionalities of the toolbox are summarized in Fig. 1. As noted previously, the main MatClassRSA code module folders, helper functions folder, and LIBSVM folder can be found in the ***src*** folder of the GitHub repository.

#### Data specifications

MatClassRSA is focused specifically on classification and RSA-adjacent analyses, and does not perform data cleaning. However, the input data specifications are aligned with the 3D space× time × trial matrix format as output by EEGLAB [22]; hence, EEGLAB is encouraged for data cleaning. Many MatClassRSA functions involve the following core inputs:

1. Input data matrix `X`: This can be a 3D space × time × trial matrix as noted above, or a 2D trial × feature matrix (for single-sensor data, for example).
2. Labels vector `Y`: This vector contains numeric stimulus labels of each trial; its length should be the size of the trial dimension of `X`.

Functions also often involve numerous additional and/or optional inputs, which are detailed in function docstrings and in the User Manual. Alternatively, some MatClassRSA functions take in outputs of other functions; for example, Visualization functions take in matrices output by Classification and RDM Construction functions.

MatClassRSA functions output variables and/or figures. Users can specify in their own code how they would like these saved in e.g.,.mat or image files.

#### Calling MatClassRSA functions

In order to use MatClassRSA, the repository needs to be in the user’s MATLAB path. Once the toolbox is in the path, specific functions are called by referencing both the MATLAB package folder name (without the ‘+’) and the specific function name. For example, to call the `noiseNormalization()` function in the `+Preprocessing` folder, the user would type the following (without the line break):

~~~
[normData, sigmaInv] =
Preprocessing.noiseNormalization(X, Y);
~~~

For succinctness, further description of MatClassRSA functions in this preprint reference only the function name, without the MATLAB package folder name prepended.

#### Main user-called functions

We now provide a narrative overview of the main user-called functions. For more information on the functions, users are encouraged to visit the User Manual in the GitHub repository (see footnote 1), which contains detailed descriptions, specifications, and example function calls. The docstrings of these main functions also contain detailed input/output specifications, and the scripts in the ***ExampleFunctionCalls*** and ***IllustrativeAnalyses*** folders of the repository illustrate various input/output specifications.

##### Module: Preprocessing

This module includes three functions that can be applied to the data prior to all other modules’ functions. First, `shuffleData()` randomizes the ordering of the trials, while keeping each trial paired with its correct label (and participant identifier, if provided). This function may be useful, for example, when the input data represent observations from multiple recordings and/or participants, with the trials ordered by recording and/or participant; the user may wish to randomize the ordering of the trials prior to calling a cross-validation classification function so that data from a given recording or participant are distributed across the cross-validation folds.

The second function, `averageTrials()` trial-averages responses to a common stimulus (and, if specified, from a common participant). As noted by Guggenmos et al. [28], these ‘pseudo-trials’ offer higher signal-to-noise ratio (SNR) than single trials, which may improve classifier performance. However, trial averaging groups of *N* trials also reduces the number of observations by roughly a factor of *N* (subject to remainder handling). Therefore, in practice, users who are interested in operating on pseudo-trials rather than single trials may want to examine tradeoffs between SNR improvements and decreased sample size when group-averaging their specific dataset.

Finally, `noiseNormalization()` accounts for variable SNR captured in each sensor and upweights or downweights the data from each sensor on the basis of its SNR. Guggenmos et al. have reported positive impacts of noise normalization on subsequent analyses such as classification [28]. However, our author team has found that at times, noise normalization can negatively impact downstream results when using MatClassRSA classification functions. Users are thus advised to examine effects of this operation on their data; more information is provided in the User Manual. This function is adapted from the code tutorial provided by Guggenmos et al. [28]^11^ and uses a helper function called `cov1para()`, created by Olivier Ledoit and Michael Wolf and imported directly from the Guggenmos tutorial into MatClassRSA.

##### Module: Reliability

Users may wish to assess the *reliability* of their data—that is, the similarity of observations across repeated instances of the data—as a complement to classification. Reliability metrics can be useful in modeling contexts because they provide an upper bound on how well *any* model will be able to explain the data. Previous versions of these functions were used for the reliability analyses reported in Figure 2 of Kong et al. [10].

**Fig. 2.**
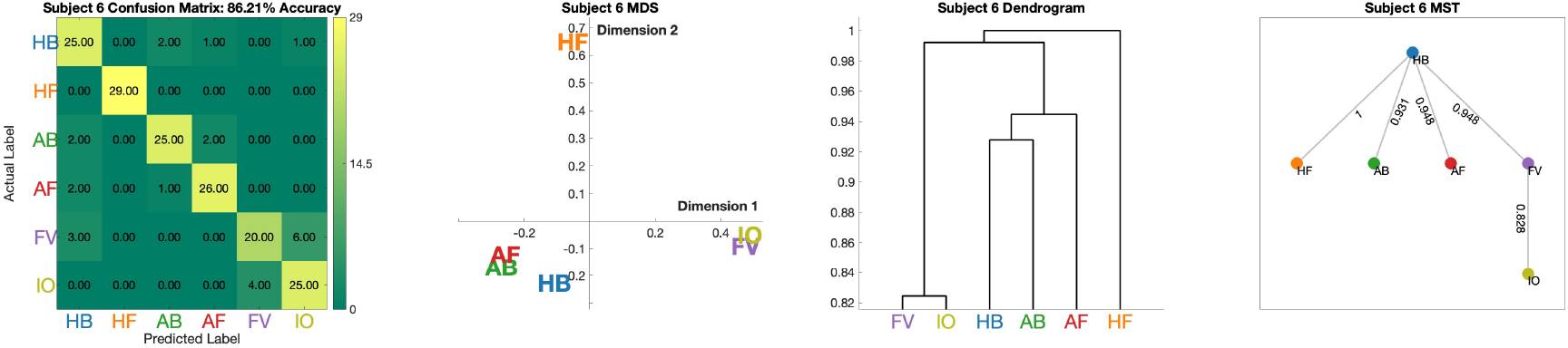
Example visualizations produced by the functions in the Visualization module. From left to right: The `plotMatrix()` function produces an image of the input matrix, which in the realm of MatClassRSA is most likely to be a multiclass confusion matrix of counts (as shown here), matrix of pairwise accuracies, or RDM; the `plotMDS()` function computes multidimensional scaling of an input RDM and then plots the coordinates of the stimuli along selected MDS dimensions; the `plotDendrogram()` function performs hierarchical clustering of an RDM and visualizes the distances among the stimuli in a tree representation; the `plotMST()` function also visualizes hierarchical clustering, this time as the paths between all stimuli for which overall distance is minimized.

The MatClassRSA Reliability module includes two functions for such assessments. Broadly, these functions involve random partitioning of the data into two split halves, trial-averaging the data on a per-stimulus basis, and then correlating the results across the stimulus dimension. These operations are repeated across several iterations, each involving a different partitioning of the data. Since splitting the data produces lower reliability estimates than if all data had been considered at once (as in e.g., a decoding analysis), the functions also apply the Spearman-Brown correction, which provides a reliability estimate across all of the data.

The `computeSpaceTimeReliability()` function computes reliabilities at each sensor and/or time point. Reliabilities can be reported as temporal trajectory for a single sensor, or a spatial distribution at a single time point; or, time- or space-averaged reliability can be reported for a given sensor or time point, respectively. The second function in this module, `computeSampleSizeReliability()`, computes reliabilities for a given sensor or time point over varying numbers of trials, which can be useful when considering the general role of sample size in classification or other RSA results.

##### Module: Classification

This module contains various functions for training and testing classifier models. Functions are separated according to multiclass and pairwise contexts. Given *N* stimulus categories, a multiclass classification is choosing among all *N* categories in model training and prediction. The output is an *N* × *N* confusion matrix; each element *i, j* expresses how many observations actually from class *i* were predicted by the model to be from class *j* (correct classifications are thus listed on the matrix diagonal, where *i* = *j*). On the other hand, pairwise classifications perform separate 2-class classifications for each pair of stimuli. Outputs from pairwise classifications are summarized in an *N* × *N* matrix, where off-diagonal elements denote the classification accuracy between classes *i* and *j*.

Functions are designed to accommodate cross validation as well as standalone training and testing (prediction) functions. Cross validation is an iterative training and testing procedure.

In *K*-fold cross-validation, the entire dataset is partitioned into *K* non-overlapping subsets called ‘folds’. Then, the classifier is trained and tested *K* times, where one fold is withheld for testing (i.e., not used for training the model) each time. Across all of the folds, each observation is used as a test observation one time. In cases where users do not wish to perform classification iteratively—for example, when some data are *only* for training or testing—MatClassRSA also offers functions that only train on the input data; these functions output models, which can be input alongside test data of the user’s choice in a predict-only function.

Finally, this module offers multiple classifiers with varying degrees of hyperparameter optimization. First, Linear Discriminant Analysis (LDA) [1] is a popular ‘out-of-the-box’ classifier that can be run without needing to tune any of its parameters. Next, Support Vector Machine (SVM) [31] contains hyperparameters that can be tuned based on attributes of the input data to further optimize classifier performance. Finally, MatClassRSA offers a Random Forest (RF) classifier [32]; this classifier also has tunable hyperparameters, though RF hyperparameter optimization is not implemented in MatClassRSA at this time.

Across these three dimensions—multiclass/pairwise, cross validation/separate train-test, and with or without hyperparameter optimization—MatClassRSA offers nine classifier functions. Cross-validation functions `crossValidateMulti()` and `crossValidateMulti_opt` handle multiclass classification, while `crossValidatePairs()` and `crossValidatePairs_opt()` handle pairwise classifications—with and without hyperparameter optimization, respectively. Separate from cross validation, `trainMulti()` and `trainMulti_opt()` handle multiclass training only while `trainPairs()` and `trainPairs_opt()` handle pairwise training only, again with and without hyperparameter optimization, respectively. These `train` functions output models, which can then be applied to new data using the `predict()` function.

In addition to the core inputs of the data matrix `X` and labels vector `Y`, users can specify subsets of data features to classify, classifier selection and—when applicable—hyperparameter optimization, feature reduction using Principal Components Analysis (PCA), data centering and scaling, and data partitioning for cross validation and/or hyperparameter optimization. When test observations have labels, statistical significance can be calculated using permutation testing—i.e., comparing the observed result to a null distribution comprising many classification attempts in which stimulus labels were randomized independently of the data trials [33, 34]. Outputs vary by function and can include the specifications of the function call; data-handling parameters related to e.g., PCA, feature subsetting, and train-dev-test partitioning; classification models; predicted labels; and p-values.

##### Module: RDM Computation

The RDM Computation module contains functions that operate both on proximity matrices (such as those output from the Classification module) and directly on the original data, such as the data matrices input to the Classification functions.

MatClassRSA Classification functions perform both multiclass and pairwise classifications. The confusion matrices output by multiclass classifications can be considered a form of similarity matrix, in that two stimuli that are confused (misclassified) more can be considered more similar than two stimuli that are never confused for one another during classification. However, as confusion matrices are count-based and also include variable values on the diagonal, they may not be optimal RDMs as is. To address this, the `computeCMRDM()` function includes options to normalize the incoming confusion matrix—for example, to represent the confusion matrix as estimated conditional probabilities [12, 35] or enforce a self-similarity measure of 1 along the diagonal [35]—as well as symmetrize, convert similarities to distances, and rank the distance values.

Similarly, the matrix of accuracies output from pairwise classifications may benefit from further transformation toward being considered an RDM. Pairwise accuracies can already be considered distance measures (higher accuracy representing better decodability and hence greater distance), but pairwise accuracies can range from 0 to 1, while chance-level classification—which would imply a distance of 0—is 0.5 [28]. Hence, the `shiftPairwiseAccuracyRDM()` function simply subtracts 0.5 from each element of an input matrix of pairwise accuracies.

The other functions in this module operate directly on the original input M/EEG data. `computeEuclideanRDM()` uses cross-validation to compute estimated pairwise distances of the original sensor-space data across the set of stimuli based on Euclidean distance. `computePearsonRDM()` similarly provides crossvalidated distance estimates, based on Pearson correlation. The cross-validation procedures applied in these functions involve multiple train/test splits of the data followed by distance or correlation measures, respectively; this approach is recommended over simply computing single distance or correlation measures across all data at once as the cross validation helps to remove noise components [28]. Both of these functions are adapted from the code tutorial provided by Guggenmos et al. [28].^12^

##### Module: Visualization

Finally, the Visualization module includes functions intended to visualize RDMs and other matrices. First, the `plotMatrix()` function creates an ‘image’ visualization of an input matrix, for example a Classification multiclass confusion matrix or pairwise accuracy, or an RDM from the RDM Computation module.

The elements of the rendered image are colored according to the corresponding value, and the function call includes options to e.g., print imaged values in the matrix elements and customize attributes of the axes and labels.

The remaining functions in this module are intended for RDM inputs specifically. First, `plotMDS()` computes Multidimensional Scaling (MDS) on the input RDM and plots coordinates of the stimulus exemplars for the specified MDS dimensions. MDS is a non-hierarchical clustering technique that converts the pairwise distances of a stimulus set to coordinates along multiple ordered dimensions [36, 37]. The stimuli can then be plotted according to their coordinates from selected dimensions—often the first two dimensions, which are the most important MDS dimensions; this can be useful for depicting clusters or separations that were not visible in the matrix view. Next, `plotDendrogram()` computes hierarchical clustering of the RDM and visualizes it as a dendrogram (tree) structure [38]. In this view, each stimulus is represented as a separate ‘leaf’ of the dendrogram, and the distance between any two stimuli is represented by the distance up the tree that one needs to traverse in order to go from one leaf to the other. Finally, `plotMST()` visualizes the distance structure of the RDM exemplars as a Minimum Spanning Tree [39]. This is another hierarchical clustering technique, whose visualization again displays all items of the stimulus set, now linked by the path that represents the minimal total distance across the set. Example visualizations produced by the functions in this module are shown in Fig. 2.

#### Helper functions

The main functions of the toolbox collectively rely on a number of helper functions. These helper functions are stored together in the `+Utils` folder. The Mat-ClassRSA User Manual includes an Appendix in which each helper function is briefly documented with function-call syntax, a narrative description, dependencies, and usage by other main and helper functions.

### Illustrative analyses

Along with the detailed function-level documentation provided in the code docstrings and in the User Manual, MatClassRSA also includes a number of illustrative analyses.

These are intended to demonstrate how multiple functions can be called together in more elaborate analyses, and also to illustrate further the functionalities of the toolbox and some of the customizable parameters of the function calls. The ***IllustrativeAnalyses*** folder of the GitHub repository contains one script per analysis documented below, which can be run by users once the example data are also downloaded.

In this section, we highlight selected results from each of the illustrative analyses provided with the toolbox. Users are encouraged to visit the User Manual in the GitHub repository (see footnote 1) to see the complete set of analyses and elaborated explanations of the analysis workflow, code snippets, and further interpretation of results.

#### Illustrative analysis 0: Download of example data files for illustrative analyses

The first illustrative analysis script does not perform an analysis per se, but rather downloads the example data files which, due to size, are not provided in the GitHub repository and are instead provided in a separate online dataset [26].

Script for this analysis

~~~
illustrative_0_downloadExampleData.m
~~~

This script locates the ***ExampleData*** folder in the user’s local MatClassRSA instance (if more than one instance is found, the first one will be used). It then iterates through (by default) all files in the online data repository [26] and downloads any that are not already found in the ***ExampleData*** folder. Users can specify a subset of files to download, or further edit the script to download files into a different directory.

The example data files are as follows:

- **Object category EEG**: ~~~
S01.mat, S04.mat, S05.matS06.matS08.mat
~~~ Single-trial EEG responses to 72 images from six visual object categories (Human Face–HF, Human Body–HB, Animal Face–AF, Animal Body–AB, Fruit/Vegetable–FV, Inanimate Object–IO). Each file contains data from a different adult participant. Each participant experienced each image 72 times in an ERP paradigm. Data files are taken directly from the OCED dataset [29]. For more information on the data and originating study, see Kaneshiro et al. [9].
- **Frequency-following responses (FFRs)**: ~~~
losorelli_100sweep_epoched.mat
~~~ Single-channel frequency-following responses (FFRs), which are auditory responses that phase lock to acoustical stimuli. Each of 13 adult participants experienced six speech and music stimuli with a fundamental frequency around 100 Hz. Each stimulus was presented 2500 times per participant. Every observation represents the average of 100 stimulus presentations for a given participant and stimulus. This data file is taken directly from the STAR-FFR-1 dataset [30]. More information on the data and the study for which they were collected can be found in Losorelli et al. [18].
- **Supporting files**: **OCEDStimuli.zip** Stimulus images for the OCED data files noted above. If the illustrative script downloads this file, it will also unzip it. These 72 images are a subset of a database of 92 images provided by Kriegeskorte et al. [15].^13^ This database, and the larger set of images from which its images are drawn, have been used in several RSA studies on visual object category processing (e.g., [8– 10, 14, 15, 40]).

##### MatClassRSA_v2_ExampleData_README.pdf

Informational document describing the dataset.

#### Illustrative analysis 1: Impact of preprocessing steps on downstream analyses

Additional data preparations, such as those provided in the Preprocessing module, can substantially impact downstream analyses. This illustrative script explores how the trial-averaging, noise-normalization, and data-shuffling functions influence characteristics of the data, multi-channel covariance, and classification outcomes.

Script for this analysis:

~~~
illustrative_1_impactOfPreprocessing.m
~~~

For example, trial averaging (using `averageTrials()`) can improve SNR of the data, while also reducing the overall voltage range of the pseudotrial averages. Fig. 3 visualizes single trials and 40-trial averaged pseudotrials of data-shuffled, noise-normalized object category data from `S01.mat` at electrode 96 (located at T4). Trial averaging makes the N170 more visually apparent reduces the overall voltage range of the data. However, trial averaging also reduces the number of available observations.

**Fig. 3.**
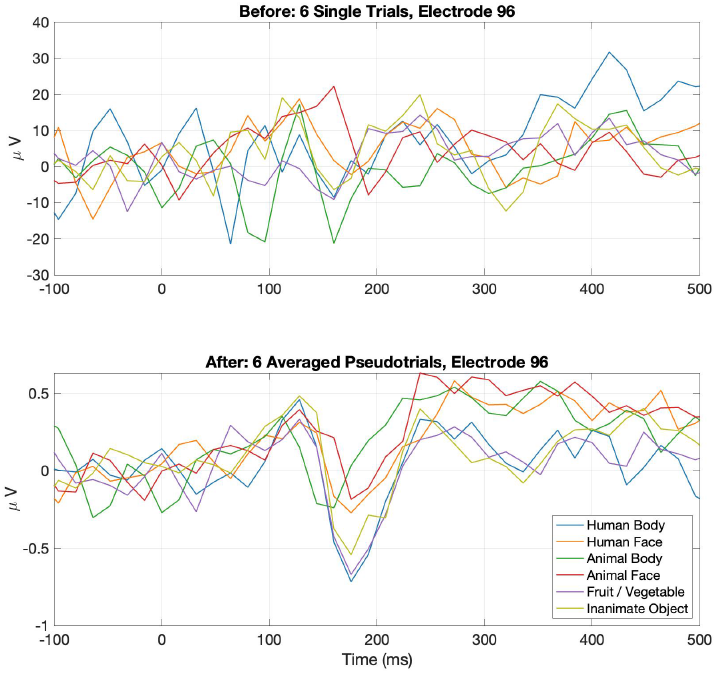
Effect of trial averaging. Visual responses to images from six object categories from `S01.mat` were averaged at a single electrode. The top plot shows single trials from each category, while the bottom plot shows averages of 40 trials per category.

Noise normalization scales each channel of data according to the inverse square root of the channel-by-channel covariance matrix. Guggenmos et al. found that this step substantially improved downstream decoding performance [28]. In the present example shown in Fig. 4, the plots on the left illustrate how calling the `noiseNormalization()` function changes the covariance structure of the `S01.mat` data. Contrary to Guggenmos et al., in the right-hand plots we find that within-participant classification performance for this specific dataset is slightly lower after noise normalization (66.47% with noise normalization in the bottom plot compared to 78.68% without in the top plot).

**Fig. 4.**
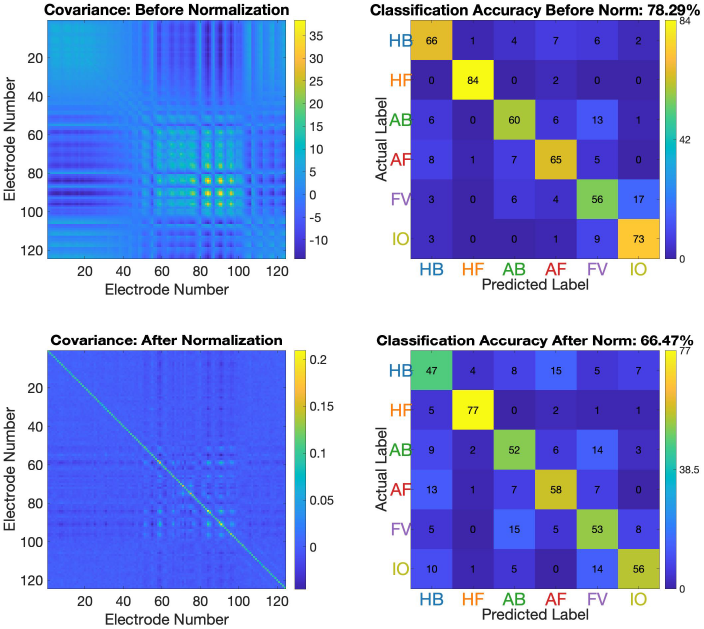
Noise normalization covariance and within-subject classification. Object category data from `S01.mat` underwent classification with and without noise normalization. The left-hand matrices illustrate the covariance matrices of the data before and after noise normalization. On the right are the 6-class classification confusion matrices. Within-subject accuracy is slightly lower after noise normalization.

Yet, when a classifier is trained on data from one participant and tested on another, we do see benefits of noise normalization. Fig. 5 illustrates an extreme case of this with SVM classifiers trained on object category data from `S01.mat` and then tested on data from `S04.mat`, `S05.mat`, and `S08.mat`. When noise normalization is not applied (top row of plots), confusion matrices from predicting on data from unseen participants show that all predictions have gravitated to the first category, resulting in classification accuracies exactly at chance level (16.67%). We surmise this is due to the classifier overfitting to training-data features that do not generalize to test data from new participants. When noise normalization is applied (bottom row of plots), the confusion matrices become somewhat more balanced, and classification accuracy improves to 25.48% and higher.

**Fig. 5.**
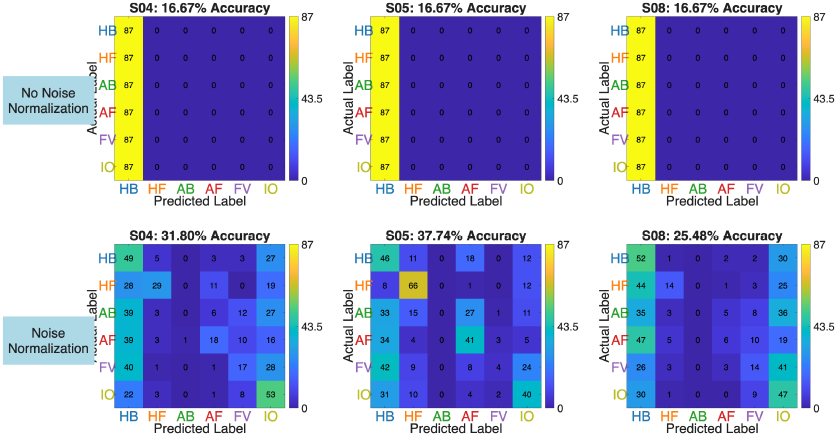
Noise normalization covariance and across-subject classification. SVM classifiers were trained on object category data from `S01.mat` and then tested on data from `S04.mat`, `S05.mat`, and `S08.mat`, with and without noise normalization. The top plots show poor classifier performance when noise normalization is not used. The bottom plots show that applying noise normalization to the data improves across-subject prediction accuracy.

Given the variable impacts of noise normalization between these within-participant and across-participant examples, users are encouraged to conduct exploratory analyses with their own data to determine whether to include noise normalization in their analysis workflow.

This script also illustrates how users can visualize their data—for example, to confirm that `shuffleData()` is working as expected. As an example, we show how the FFR dataset—which includes both stimulus labels and optional participant labels for each trial—can be passed into this function with the optional specification to keep participant identifiers coupled with the data observations and labels. Fig. 6 shows the category labels of the ordered trials, before and after data shuffling. As shown in the top plot, the trials as initially loaded are ordered by stimulus: All trials from category 1 followed by all trials from category 2, and so on. If this trial ordering were carried into cross validation, certain classes would be over- or under-represented in each cross-validation partition; as an extreme example, performing 6-fold cross-validation on this ordering of the trials would equate to partitioning the data exactly by class, ensuring that the class of each holdout test partition is not represented *at all* in the training partitions! The bottom plot of this figure shows that after data shuffling, the six categories are well distributed throughout the ordered trials, ensuring a good balance of classes across future data partitions. Fig. 7 depicts the per-participant averaged data across all stimuli before and after data shuffling (each line represents a participant), providing visual confirmation that the mappings between data trials and participant labels are preserved during data shuffling.

**Fig. 6.**
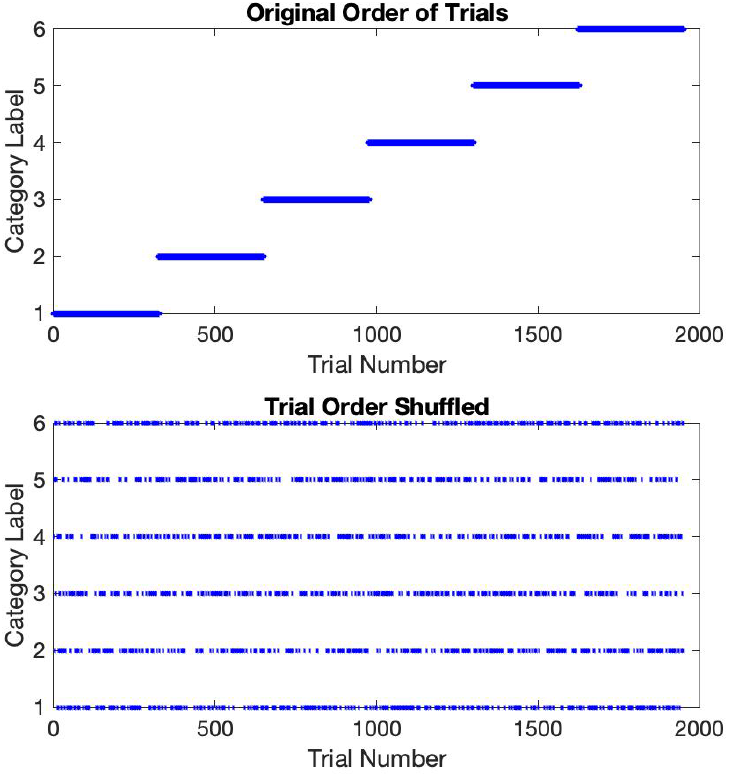
Distribution of trials before and after data shuffling. The example data file from the Losorelli et al. dataset [30] initially has trials grouped by stimulus. Shuffling the data ensures a better distribution of the stimuli across the dataset.

**Fig. 7.**
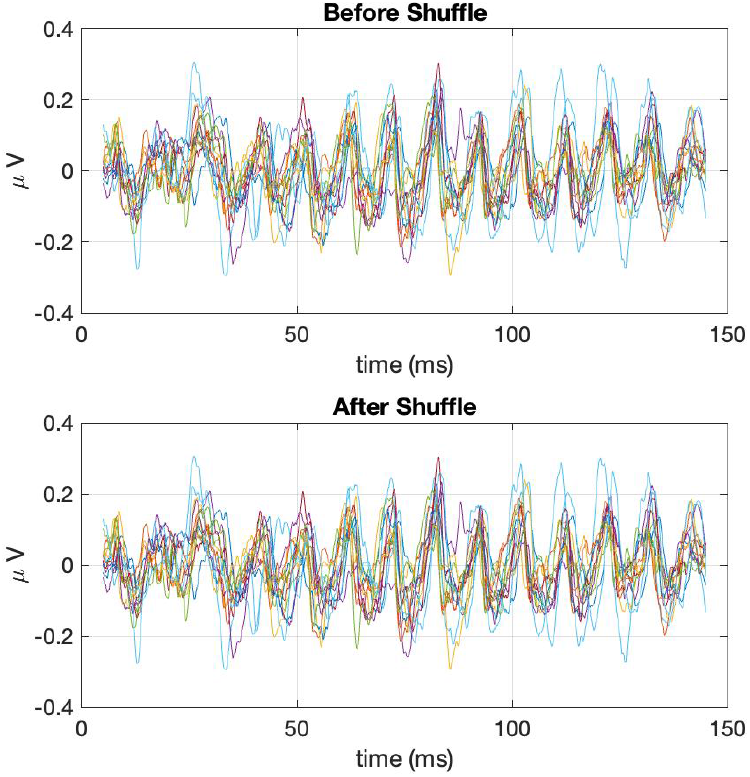
Per-participant averaged data are consistent before and after data shuffling. FFR responses phase locked to acoustical stimuli with a fundamental frequency of 100 Hz were input to the `shuffleData()` function with the optional participant identifier included. Each line represents the average of all data (across all stimuli) for one participant before and after averaging. This figure provides visual confirmation that the function preserved the data-participant mappings.

Additional sub-analyses, figures, and interpretations for this illustrative analysis can be found in the User Manual.

#### Illustrative analysis 2: Single-channel analyses

Multi-channel M/EEG data offer rich spatiotemporal information. As an alternative to visualizing spatial weight vectors output by classifiers—which can lack visual smoothness (e.g., [13]) and require special care in interpretation [41]—single-channel analyses can offer broad insight into relevant spatial features.

Script for this analysis:

~~~
illustrative_2_singleChannelAnalyses.m
~~~

For example, users may wish to identify electrodes that record reliable data. In Fig. 8, single-channel reliabilities for object category participant `S01.mat`, computed using `computeSpaceTimeReliability()`, are displayed on a scalp map, highlighting especially reliable data recorded from electrodes over right occipital cortex.^14^

**Fig. 8.**
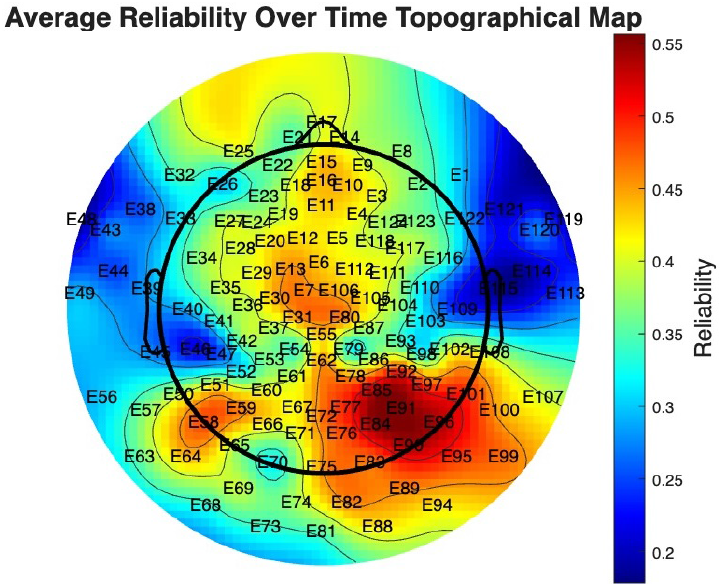
Topomap of single-channel reliabilities. Average reliability across all time points was computed for each sensor of the `S01.mat` object category data and visualized on a scalp map. Sensors over right occipital cortex recorded the most-reliable data.

While reliability analyses can help to identify channels that record consistent data over repeated trials, single-channel classification analyses can be a useful complementary analysis to derive scalp maps highlighting sensor locations whose data classify well and may therefore be task-relevant. An example from the same participant, with data classified using LDA with the `crossValidateMulti()` function, is shown in Fig. 9. We see broad alignment between this scalp map and the reliability map, as well as with the group-averaged results originally reported by Kaneshiro et al. [9].

**Fig. 9.**
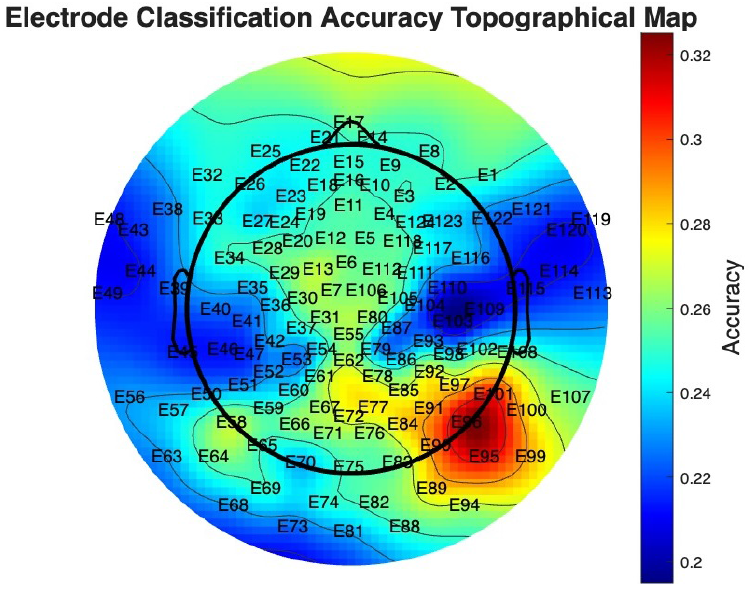
Topomap of single-channel classification rates. Six-class classifications of the object category data from `S01.mat` were performed independently on data from each electrode. Similar to the reliabilties map, the topomap of classifier accuracies highlights right occipital cortex.

Further elaboration, figures, and interpretations for this analysis are described in the User Manual.

#### Illustrative analysis 3: Time-resolved analyses

One of the greatest assets of M/EEG data is high temporal resolution. While multivariate analyses such as classification can optimize high-dimensional data, restricting input data to pre-identified relevant temporal features can aid data interpretation, reduce processing time, and even boost classifier performance. MatClassRSA functions can be used to analyze data subsets along the time dimension.

Script for this analysis:

~~~
illustrative_3_timeResolvedAnalyses.m
~~~

First, as a complement to single-channel reliabilities, the `computeSpaceTimeReliability()` function can also be used to compute reliabilities across all electrodes at each time point; the results can then be plotted as time course. Fig. 10 shows the reliability over time of object category participant `S01.mat`, computed at the exemplar (72-class) level across all electrodes. As can be seen from the plot, reliability remains near zero in the pre-stimulus and early post-stimulus intervals until around 80 msec post-stimulus onset—around the time of the P1 component. It then peaks around the timing of the N170 component and decreases toward the end of the response epoch.

**Fig. 10.**
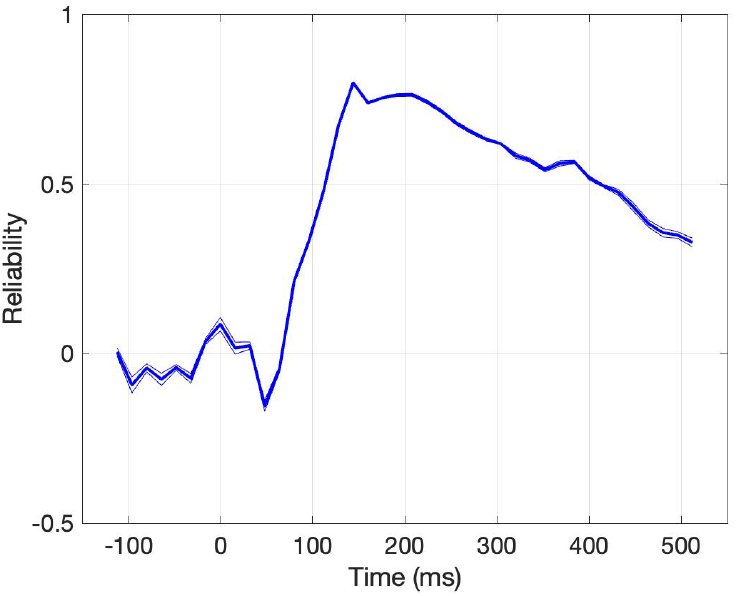
Time-resolved reliability, averaged across electrodes. Reliability, averaged across all electrodes, was computed for each time point of `S01.mat` data and visualized. Non-negative reliabilities correspond in time to the P1 ERP component, and peak around the time of the N170 component.

This type of temporal information can be useful in specifying classification analyses as well. Fig. 11 involves the classification of shuffled, group-averaged data (15 trials per group) from object category data from `S01.mat` using `crossValidateMulti()` with LDA. Here, when data from all time points (from -112 to 512 msec relative to stimulus onset) and electrodes are included for 6-class classification, the classifier accuracy of 42.69% is well above chance level of 16.67%. However, when data from only 128 to 208 msec post-stimulus onset are input to the classifier, the classifier accuracy improves to 69.88%, with a stronger confusion matrix diagonal.

**Fig. 11.**
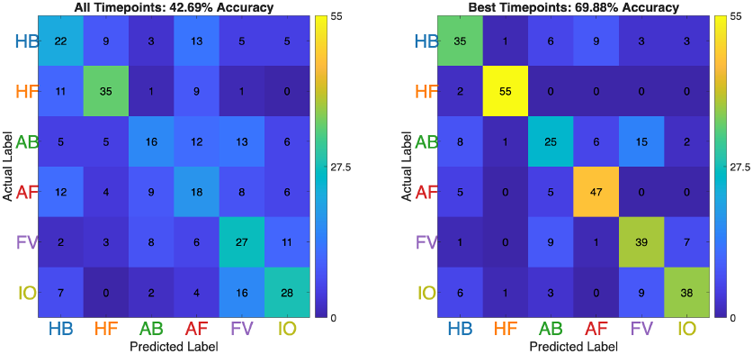
Confusion matrices when classifying over all time and best time window. Six-class classifications of the object category data from `S01.mat` were performed across the entire response epoch (-112 to 512 msec—left) as well as across the best time points (128 to 208 msec—right) as identified by a sliding-time-window classification (described in the User Manual). Compared to classifying all available data, restricting the time dimension of the input data to the best-performing segment improves classification performance from 42.69% to 69.88%.

Additional reliability, ERP, and classification results as well as visualizations are described in detail in the User Manual.

#### Illustrative analysis 4: Train-test SVM optimization

MatClassRSA classification functions with optimization offer fairly broad default grids over which to tune the SVM hyperparameter(s). In this illustrative analysis, we demonstrate how SVM hyperparameters can be tuned over successive grid searches, where optimal parameters from one attempt can specify a finer grid in the next, and further improve classifier performance over hyperparameters optimized in the default grid specification only.

Script for this analysis:

~~~
illustrative_4_trainTestOptimizeSVM.m
~~~

To begin, we performed a 6-class SVM classification on the object category data from `S01.mat`. We called the `crossValidateMulti_opt()` function from the Classification module, using the rbf kernel and optimizing *γ* and `C` using default grid parameters (5 logarithmically spaced points between 10^−5^ and 10^5^). As shown in Fig. 12, classifier performance was 66.48%, well above a chance level of 16.67%.

**Fig. 12.**
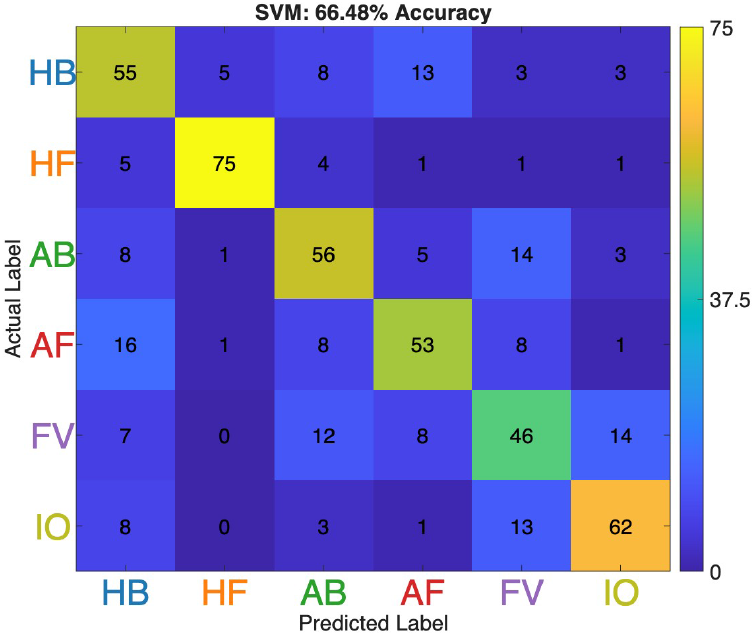
Confusion matrix for SVM classification with default grid-search hyperparameters. Six-class classifications of the object category data from `S01.mat` were performed using SVM with an rbf kernel. Hyperparameters *γ* and `C` were optimized over the default grid parameters.

The optimal hyperparameters identified in the first classification’s default grid search were *γ* of 0.0032 and `C` of 316. In a follow-up classification, we specified finer-grained *γ* and `C` grid parameters around these initial optimal values. This second round of optimization identified optimal gamma of 0.0146 and optimal C of 1000, and improved classifier performance to an accuracy of 81.23% as shown in Fig. 13.

**Fig. 13.**
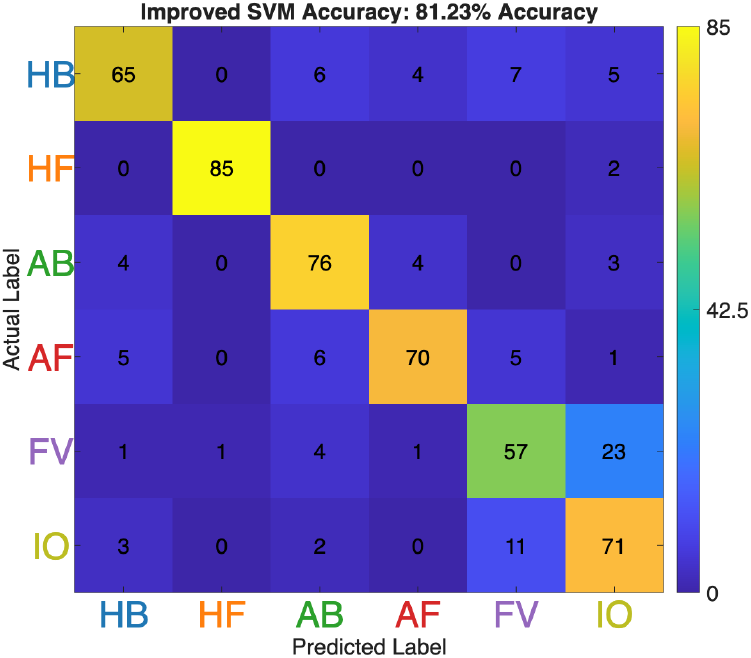
Confusion matrix for SVM classification with refined grid-search hyperparameters. Following from the previous figure, when a second SVM rbf classification is performed on `S01.mat` data with hyperparameters optimized on a finer grid based on the optimal `γ` and `C` from the initial optimization, the overall accuracy improves.

Further elaboration and interpretations for this analysis are described in the User Manual.

#### Illustrative analysis 5: Comparison of different RDM constructions

MatClassRSA offers various means for creating RDMs, both from classifier outputs and directly from the original sensor-space data. This analysis provides examples of different RDM constructions.

Script for this analysis:

~~~
illustrative_5_compareRDMs.m
~~~

We first performed both multiclass and pairwise 6-class classification of `S06.mat` object category data using LDA with the `crossValidateMulti()` and `crossValidatePairs()` functions, respectively. Classifier outputs were converted to RDMs. RDMs were percentile ranked for visualization only and plotted using the `plotMatrix()` function, while MDS and dendrogram plots were created from non-percentile-ranked RDMs and visualized using the `plotMDS()` and `plotDendrogram()` functions. As shown in Fig. 14, the percentile-ranked RDMs look similar across the two approaches. The MDS and dendrogram plots also look similar in overall shape, although the distance values are larger for the multiclass approach compared to the pairwise.

**Fig. 14.**
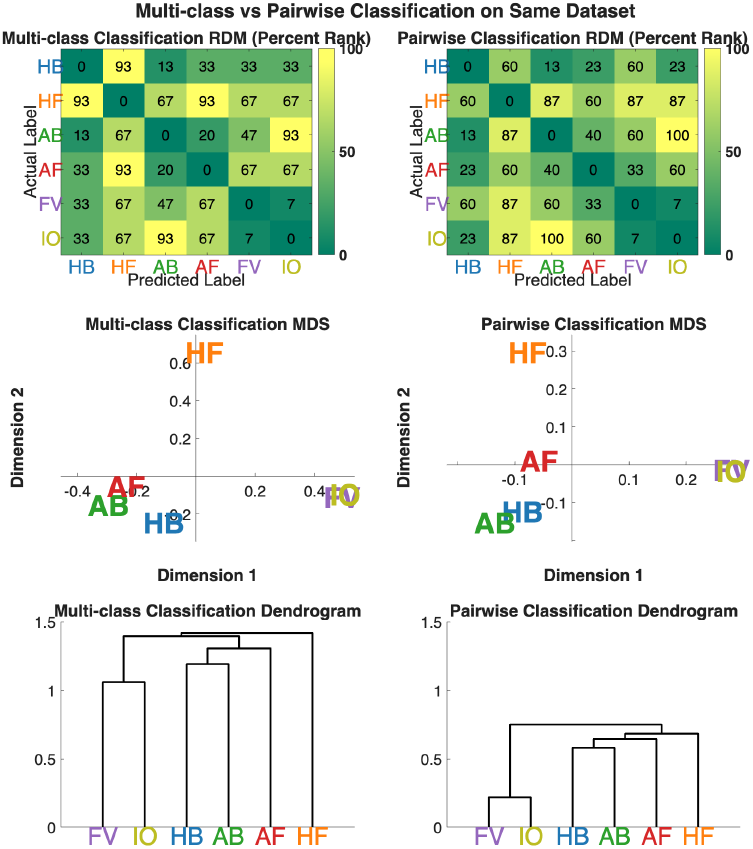
Six-class multiclass and pairwise accuracy visualizations. Object category data from `S06.mat` underwent 6-class classifications. Top: Percentile-ranked RDMs have a highly similar distance structure. Middle and bottom: Visualizations of RDMs (not percentile ranked) as MDS and dendrograms show similar category structure across the two approaches, with larger values for multiclass.

We also computed 6-class Pearson and Euclidean RDMs directly from channel 96 of the `S06.mat` data. Data underwent shuffling, noise normalization, and trial averaging prior to being input to the `computePearsonRDM()` and `computeEudlideanRDM()` functions. The RDMs were then visualized using both the `plotMatrix()` and `plotMST()` functions. Fig. 15 shows the resulting RDMs (top), which have been percentile-ranked for visualization. The middle and bottom rows show multidimensional scaling plots and minimum spanning trees, respectively, with distances reported in their original units (note that MST path lengths do not convey relative path distance values). Even with differing ranges of distance values for each RDM technique as shown in MDS and MST plots, the Pearson and Euclidean approaches produce similar arrangements of the stimulus categories.

**Fig. 15.**
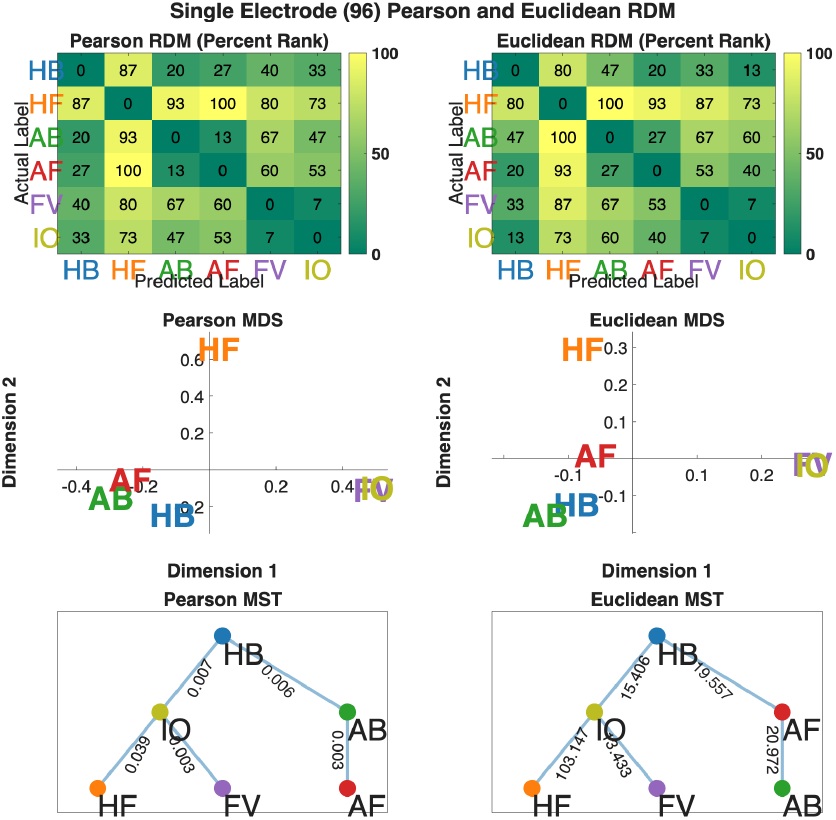
Six-class Pearson and Euclidean RDMs, MDS, and MST plots. Six-class RDMs were computed from `S06.mat` data, channel 96. Top: Percentile-ranked RDMs are broadly similar across the two techniques. Middle: Stimulus categories show similar groupings along the first two MDS dimensions. Bottom: MST plots are also highly similar in structure, even while the specific (non-percentile-ranked) distances occupy different data scales across the two techniques.

Finally, we compared 72-class Pearson and multiclass classification RDMs and MDS computed from channel 96 of the same `S06.mat` data. Results are summarized in Fig. 16. Contrary to previous 6-class figures, we see more variability between the two approaches: The structure of certain categories is more visually evident in the Pearson than the classification RDM, reflected in blocking along the RDM diagonal particularly for Human Face exemplars (rows 13–24) and Animal Body and Face exemplars (rows 25–48). Human Face and Animal Face exemplars also appear to form tighter clusters in the Pearson MDS plot, as shown by the orange and red dots, respectively. For both MDS plots, Human Face exemplars (orange dots) appear to be relatively well clustered and separated from other categories.

**Fig. 16.**
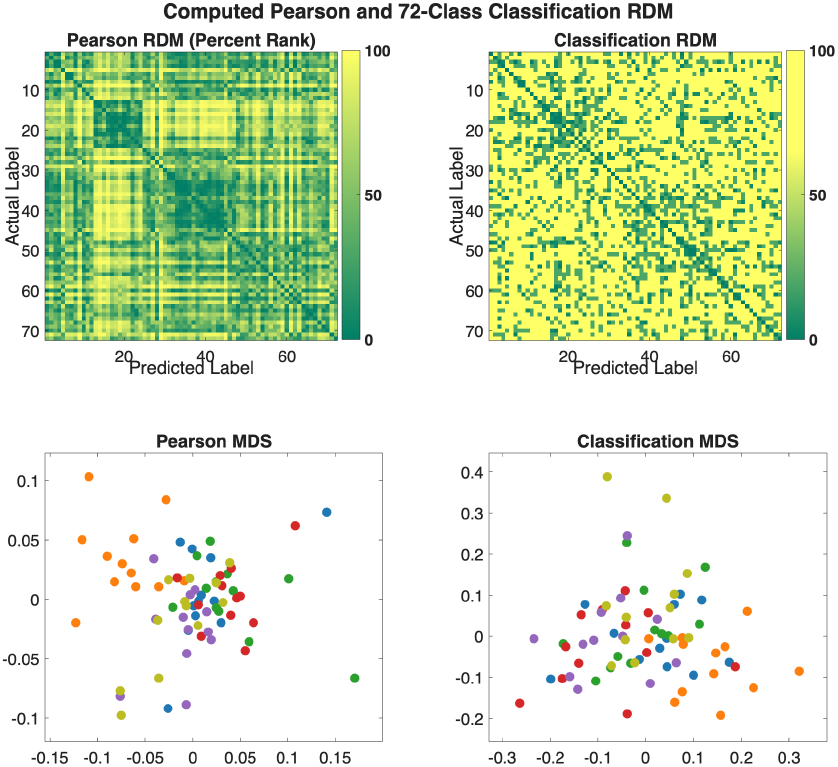
72-class Pearson and classification-based RDMs. Channel 96 of the `S06.mat` object category data were used to construct 72-class Pearson and multiclass classification RDMs. These approaches produce markedly different results: By inspection, the Pearson RDM (top) shows a clearer category structure in e.g., rows 13–24 (Human Face) and rows 25–48 (Animal Body/Face) as well as stronger clustering of some categories (colors) in the MDS plot (bottom). Orange dots, which reflect Human Face exemplars, appear to separate from other categories for both RDM construction approaches.

For more information, figures, and interpretations for these analyses, please visit the MatClassRSA User Manual.

#### Illustrative analysis 6: Customizing figures with stimulus images

The final illustrative analysis demonstrates how the basic MatClassRSA visualization functionalities can be customized and extended—for example to annotate plots with images rather than text labels.

Script for this analysis:

~~~
illustrative_6_figureCustomizations.m
~~~

Trial-averaged data from object category dataset `S01.mat` underwent 6-class classification. In the confusion matrix shown in Fig. 17, rows and columns are labeled with example images from each stimulus category rather than the usual numeric or text labels. Users can leverage this example to customize stimulus labels in their own matrix, MDS, dendrogram, and MST plots.

**Fig. 17.**
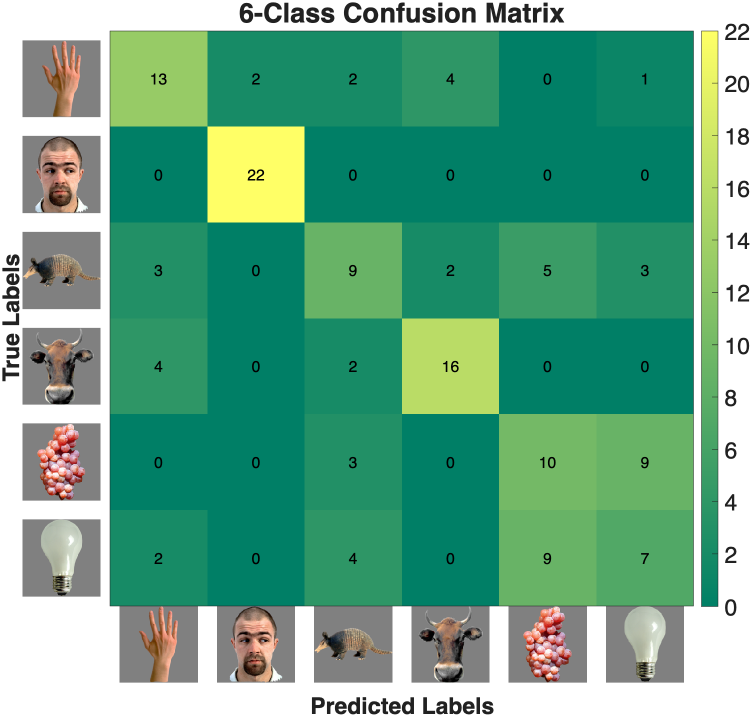
6-class confusion matrix customized with stimulus images. The object category data from `S01.mat` underwent 6-class classification. The confusion matrix visualization has been customized with an example image from each of the stimulus categories. The images shown in this figure are from the database of object category images provided by Kriegeskorte et al. [15]; see the description of Illustrative Analysis 0 and footnote 13, above, for more information.

Further elaboration and interpretations for this analysis are provided in the User Manual.

## Discussion

We have presented the v2 release of MatClassRSA, a MATLAB toolbox that classifies M/EEG data, performs additional preprocessing and reliability analyses, and also converts M/EEG data and classifier outputs to RDMs and visualizes them. The current release offers a restructured and expanded codebase as well as an updated User Manual and illustrative analyses. Numerous studies have demonstrated the utility of M/EEG classification as a data-driven tool for analyzing high-dimensional data and data with large stimulus sets, and as a means of deriving RDMs that can then be compared with other representations of the stimulus set using RSA. Classifying spatial and/or temporal subsets of the input data in ‘searchlight’ approaches also provides a data-driven means of identifying task-relevant features of the neural data. The predictive nature of classification in general also highlights potential diagnostic and clinical applications.

MatClassRSA is by no means the only resource for M/EEG classification; a number of other MATLAB toolboxes— such as BCILAB [43], CoSMoMVPA [44], ADAM [45], EEGsig [46], and MVPAlab [47]—as well as Python toolboxes with M/EEG classification functionalities [48–52], decoding toolboxes for other neuroimaging data modalities [53, 54], and more general analytic frameworks for M/EEG [22, 55–57] are freely available, along with tutorials (e.g., [28, 58, 59]). As part of the growing ecosystem of M/EEG decoding toolboxes, MatClassRSA focuses on constructing and visualizing RDMs using classification and other approaches. The expanded offerings of the v2 release—including preprocessing steps [28], reliability calculations, and non-classification RDM constructions— complement the functionalities of other toolboxes. Users are encouraged to explore their options to find the best tool to suit their analysis and workflow needs.

We acknowledge limitations of the current work. For example, the selection of classifiers remains fairly limited; and the toolbox is currently not designed to handle data with missing values, nor is it validated for data with highly imbalanced classes. Interpretation and visualization of classifier weights is not currently implemented, and statistical analyses are limited to permutation testing for single classification runs. By design, MatClassRSA does not include upstream data cleaning or downstream RSA analysis functionalities, and users are encouraged to engage other toolboxes already specializing in those steps [22, 23]. Finally, the lack of a GUI may hinder adoption of MatClassRSA by researchers who are accustomed to working in graphical environments. However, we hope that the extensive documentation and example code provided in the toolbox repository lowers the barrier to entry for *all* researchers, while arguably promoting better reproducibility and shareability of analyses through script-and function-based implementations [44].

Future work on MatClassRSA can involve implementing additional functionalities within the toolbox, or providing bridging functionalities to better connect MatClassRSA function outputs to complementary functions of other toolboxes, such as the creation of ‘publication-ready’ figures [45] and advanced statistics [44, 45, 47]. While the toolbox has been validated with multiple datasets, we expect that real-world usage with new data will bring to light new bugs, edge cases, and feature requests that can be addressed in future versions. Finally, we will be assessing usability of the current release with researchers from various disciplines including Education, Psychology, and Computer Science, to identify how the toolbox can be improved with regard to its implementation and documentation.

## ACKNOWLEDGEMENTS

This research was supported by the Patrick Suppes Gift Fund and the Roberta Bowman Denning Fund for Humanities and Technology (BK). The authors gratefully acknowledge Yiran Duan, Steven Losorelli, Vaidehi Natu, Duc T. Nguyen, Ethan Roy, Greta Vilidaite, and Xiaoqian Yan for their valuable feedback and insights contributed while using development versions of the toolbox. We also thank Peter J. Kohler for technical suggestions, Daniel O’Leary and Andero Uusberg for advice on input data specifications, Amilcar Malave for guidance on the code release, and an anonymous reviewer for constructive feedback on a previous version of the toolbox. Finally, we thank Marcos Perreau Guimaraes and Hyung-Suk Kim for their past contributions that are reconceptualized in this toolbox; and Patrick Suppes for championing M/EEG classification in cognitive neuroscience research.

This preprint was prepared using the *HenriquesLab bioRxiv* Overleaf template by Ricardo Henriques.^15^

## AUTHOR CONTRIBUTIONS

Designed the toolbox architecture: BCW, AMN, BK. Implemented the code: BCW, RG, NCLK, BK. Validated the code: BCW, RG, NCLK, BK. Documented the toolbox: BCW, RG, NCLK, BK. Created illustrative analyses: RG, BK. Provided theoretical and domain expertise: FR, AMN. Wrote the preprint: BK. Provided feedback and editing: BCW, RG, NCLK, FR, AMN, BK.

## DECLARATION ON THE USAGE OF AI

The authors used Azure OpenAI gpt-4o-mini through Stanford University’s AI Playground platform^16^ for assistance in performing Git and L_A_TEX operations as well as for clarifying, debugging, and documenting code files written by co-authors who had become less active on the project. We did not otherwise use any AI tools for developing the code or for writing the user manual or preprint.

## Footnotes

1 https://github.com/berneezy3/MatClassRSA

2 https://zenodo.org/records/17850472

3 https://purl.stanford.edu/kv831rr3606

4 https://choosealicense.com/licenses/mit/

5 https://www.mathworks.com/products/matlab.html

6 https://www.mathworks.com/products/parallel-computing.html

7 https://www.mathworks.com/products/statistics.html

8 https://www.mathworks.com/products/image-processing.html

9 http://www.csie.ntu.edu.tw/~cjlin/libsvm

10 https://github.com/m-guggenmos/megmvpa/blob/master/tutorial_matlab/matlab_distance.ipynb

11 https://github.com/m-guggenmos/megmvpa/blob/master/tutorial_matlab/matlab_distance.ipynb

12 https://github.com/m-guggenmos/megmvpa/blob/master/tutorial_matlab/matlab_distance.ipynb

13 https://bit.ly/3Ma1ZSZ, accessed November 19, 2025.

14 The function to render scalp maps—developed as a standalone function by the Parra Lab (https://parralab.org/) and freely available on GitHub [42] (https://github.com/dmochow/SRC/blob/master/topoplot_new.m)—is included in the MatClassRSA +Utils folder with attribution.

15 https://www.overleaf.com/latex/templates/henriqueslab-biorxiv-template/nyprsybwffws

16 https://uit.stanford.edu/service/aiplayground

